# Phase variation of *Clostridioides difficile* colony morphology occurs via modulation of cell division

**DOI:** 10.1101/2025.08.20.671228

**Authors:** Anchal Mehra, Elizabeth M. Garrett, Christopher J. Serody, Rita Tamayo

## Abstract

Phase variation of *C. difficile* colony morphology occurs via modulation of transcription *cmrRST*, which encodes a three-protein signal transduction system. Response regulators CmrR and CmrT promote rough colony development, cell elongation and chaining, surface motility, and disease in the hamster model of infection, while impairing swimming motility and biofilm formation. Using RNA-Seq, we identified the CmrR and CmrT-dependent transcriptional differences in rough and smooth colonies. Further analysis showed that CmrT, but not CmrR, is required for differential expression of most of the genes. Two CmrT-regulated genes, herein named *mrpA* and *mrpB*, were together sufficient for restoring all CmrT-dependent *in vitro* phenotypes in a *cmrT* mutant and alleviating selection of *cmr* phase ON cells during growth on an agar surface. MrpA and MrpB are uncharacterized proteins with no known function but are highly conserved in *C. difficile*. Using immunoprecipitation and mass spectrometry to identify interacting partners, we found that MrpA interacts with the septum-site determining protein MinD and several other proteins involved in cell division and cell shape determination. Ectopic expression of *mrpAB* resulted in atypical cell division, consistent with MrpAB interference with MinD function. Our findings reveal a potential mechanism by which phase variation of CmrRST modulates colony morphology and motility: in *cmr* phase ON cells, CmrT-mediated expression of *mrpAB* interferes with normal cell division resulting elongated cells that enable expansion of the population across a surface while limiting swimming motility.

**AUTHOR SUMMARY:** *C. difficile* can reversibly switch between two colony morphology variants that differ in multiple additional phenotypes. Previous work determined that the phenotypic switch occurs through phase variation of the CmrRST signal transduction system, however the CmrRST-regulated genes that mediate these phenotypes were unknown. Here, we identified the CmrR- and CmrT-regulated genes that are differentially expressed between rough and smooth colonies and identified two genes, *mrpA* and *mrpB*, that together are sufficient to confer phenotypes associated with *cmrRST* expression. Protein interaction studies revealed that MrpA interacts with MinD, a protein that helps ensure symmetric cell division in bacteria. Based on these findings, we propose that MrpA disrupts MinD function leading to aberrant cell division, resulting in elongated cells that form rough colonies. This study reveals previously uncharacterized proteins that affect *C. difficile* cell division and broadly impact the physiology of this pathogen.

## INTRODUCTION

Many bacterial species generate phenotypic heterogeneity to increase the fitness of a population as a whole. Some species benefit from phenotypic heterogeneity due to the division of labor among the distinct subpopulations that specialize in performing different tasks. Phenotypic diversity may also serve as a bet-hedging strategy, increasing the chances of a subpopulation surviving an environmental stress (1).

*Clostridioides difficile* is an intestinal pathogen with a significant global health burden, causing disease ranging from mild diarrhea to pseudomembranous colitis and sepsis. Many *C. difficile* strains display colony dimorphism—on an agar surface, colonies appear rough with filamentous edges or smooth with rounded edges (2–4). Prior work has established that *C. difficile* colony morphology is subject to phase variation, which generates a phenotypically heterogeneous population through stochastic, reversible ON/OFF modulation of gene expression (3, 5). Several other phenotypes are associated with each colony morphology. The rough colony variant forms elongated, chained cells and exhibits greater surface motility, while the smooth colony variant produces more biofilm and shows more swimming motility (3). The rough colony variant also trends toward greater virulence than the smooth colony variant in the hamster model of acute *C. difficile* infection despite comparable toxin production.

Phase variation in *C. difficile* can occur through site-specific recombination that reversibly inverts regulatory DNA elements (5, 6). One of these elements, the *cmr* switch, controls colony dimorphism and the associated phenotypes (3). The *cmr* switch contains a promoter that, when oriented toward *cmrRST*, drives transcription of the operon and leads to the *cmr*-ON state and rough colony development; when the *cmr* switch is in the inverse orientation, this transcription is lost, leading to the *cmr*-OFF state and smooth colonies (**Fig. 1A**) (3, 7).

**Figure 1.**
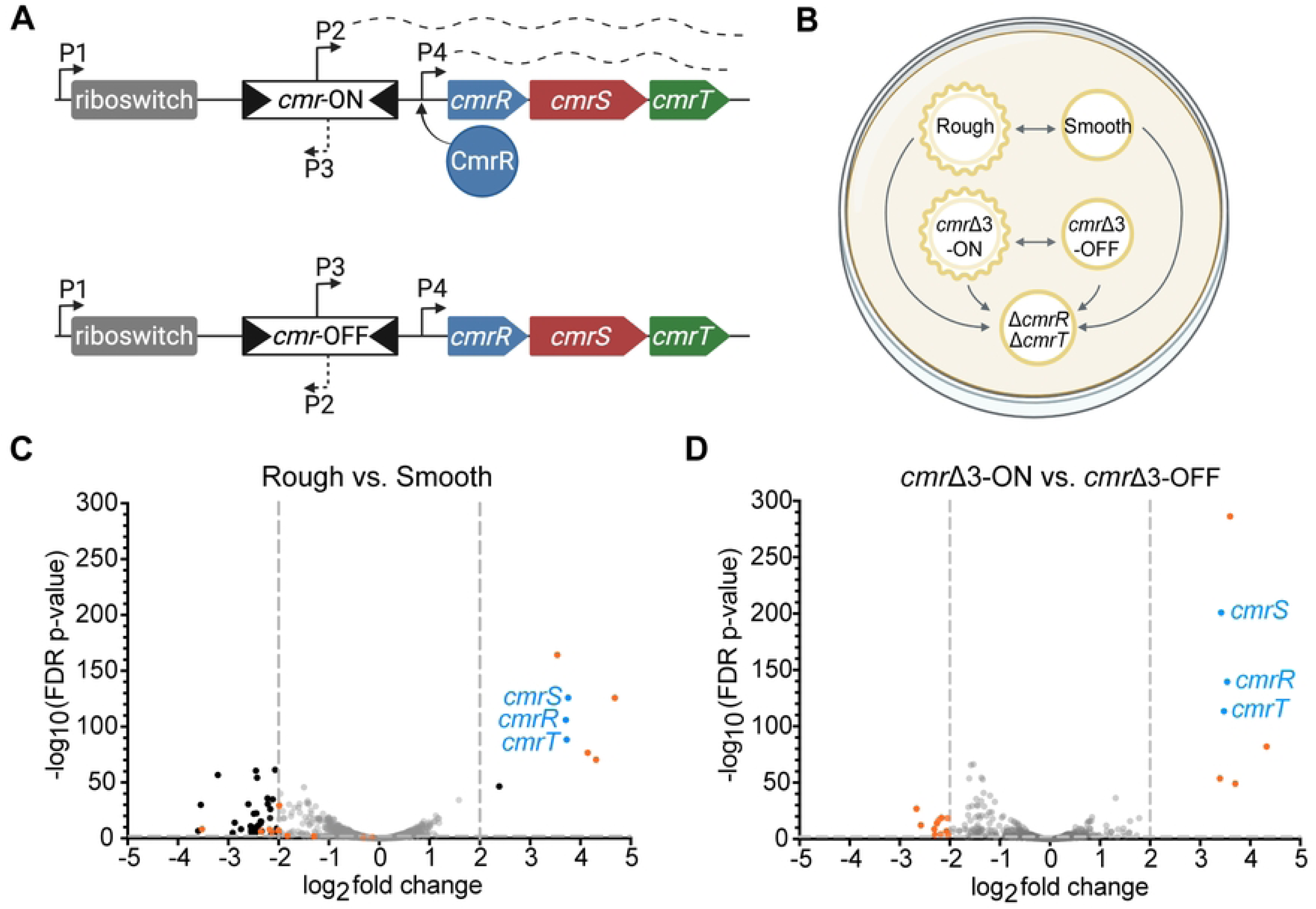
RNA-Seq reveals an overlap in gene expression in wildtype rough *cmr*Δ3-ON colonies. **(A)** Diagram of the *cmrRST* operon with the ON or OFF *cmr* switch orientation (top and bottom, respectively). Arrows represent promoters; dashed lines represent transcription. The position of the c-di-GMP riboswitch that positively regulates expression is indicated. **(B)** Overview of study design. Wildtype rough, wildtype smooth, *cmr*Δ3-ON, *cmr*Δ3-OFF, and Δ*cmrR*Δ*cmrT* strains were grown on BHIS-agar for 24 hours before collection of growth for RNA-Seq analysis. Arrows show the comparisons made. **(C-D)** Volcano plots of differences in transcript abundance between **(C)** wildtype rough versus smooth colonies and **(D)** *cmr*Δ3-ON vs. *cmr*Δ3-OFF mutants. Gray dotted lines demarcate cutoffs of > 2 log_2_ fold-change and/or FDR *p*-value < 0.01. Black dots indicate genes that met the cut-offs; gray dots indicate those that did not. Blue dots denote *cmrRST*. Orange dots highlight genes that were differentially expressed in the *cmr*Δ3-ON versus -OFF dataset and that met cutoffs.

The *cmrRST* operon encodes a signal transduction system: CmrS is a predicted histidine kinase for which the activating signal is unknown, and CmrR and CmrT are annotated as OmpR-family response regulators (8, 9). CmrR and CmrT each have an N-terminal phosphoreceiver domain and a C-terminal winged helix DNA binding domain, suggesting that they serve as transcriptional regulators (5, 10). These regulators appear to serve discrete roles. CmrR is required for intestinal colonization in the hamster model of infection, decreases biofilm production, and autoregulates the *cmrRST* operon, whereas CmrT is required for rough colony development, surface motility, cell chaining, and cell elongation (3, 7). Despite the broad effects of CmrRST on *C. difficile* physiology, the genes regulated by CmrR and/or CmrT that mediate *cmr-* associated phenotypes have not been identified. CmrRST may regulate a singular pathway that coordinately modulates all the *cmr*-associated phenotypes, or it may control multiple genes, each responsible for one or a subset of the phenotypes.

In this study, we used RNA-Seq to define the CmrR and CmrT regulons and their contributions to the rough and smooth colony transcriptomes. We identified two CmrRST-regulated genes, which we named <u>m</u>odulators of rough <u>p</u>henotype A and B (*mrpA* and *mrpB*), that when co-expressed promote rough colony formation, surface motility, and cell elongation and chaining while impeding swimming motility. We found that MrpA interacts with septum-site determining protein MinD (11–13). Consistent with this observation, expression of *mrpAB* induces abnormal cell division, with misplaced septa and elongated cells. Based on our results, we propose that phase variation of CmrRST, and thereby MrpAB, modulates cell division in *C. difficile*, with the *cmr*-ON state resulting in MrpAB-mediated disruption of proper cell division. The consequent elongated, chained cells may underlie the enhanced surface motility and reduced swimming motility of the rough colony variant.

## RESULTS

### CmrRST accounts for some, but not all, transcriptional differences between rough and smooth colony variants

To determine the transcriptional differences that may drive colony dimorphism in *C. difficile*, we compared the transcriptomes of five strains grown on an agar surface. Specifically, rough and smooth colony isolates of wildtype R20291 were grown alongside phase-locked mutants, which were previously made by deleting three nucleotides in the right inverted repeat to prevent *cmr* switch inversion (7). These *cmr*Δ3-ON and *cmr*Δ3-OFF mutants form exclusively rough and smooth colonies, respectively. Because *cmrRST* transcription is controlled by multiple promoters (**Fig 1A**) (7), a Δ*cmrR*Δ*cmrT* mutant that forms only smooth colonies was also included. **Fig. 1B** shows the five strains and the RNA-Seq comparisons conducted to identify differentially expressed genes (DEGs) that met cutoffs of log_2_ fold change > 2 and FDR adjusted *p*-value < 0.01. Locus numbers formatted as CDR20291_XXXX in the R20291 genome (GenBank: FN545816.1) are abbreviated CDRXXX below.

RNA-Seq analysis of the wildtype rough versus smooth colony variants revealed 49 DEGs (**Fig. 1C**, **Table S3**), 41 of which were more highly expressed in smooth colonies relative to rough colonies. Fifteen of these 41 genes are predicted to have phage-related functions, and ten are uncharacterized or hypothetical. Several metabolic genes had higher expression in smooth colonies including genes involved in the synthesis of riboflavin (CDR1595), and the utilization of ethanolamine (CDR1833-1838), mannitol (CDR2221-2222), or pyruvate (CDR2301). The remaining genes more highly expressed in smooth colonies have varying functions, including signaling, transport, and a protease. Of the eight genes more highly expressed in rough colonies, three have no predictive annotations (CDR1689, CDR1690, and CDR1914), one encodes the cell wall protein Cwp28 (CDR1911) (14, 15), and the remaining four are involved in signaling (*cmrRST* and the c-di-GMP phosphodiesterase gene CDR2040) (16, 17). That the *cmrRST* operon is among the most abundant transcripts in rough colonies is consistent with prior work showing greater *cmrRST* transcript in rough colonies than in smooth (3, 7).

The *cmr*Δ3-ON and *cmr*Δ3-OFF transcriptomes had 18 DEGs (**Fig. 1D, Table S3**). All seven genes that had higher expression in the *cmr*Δ3-ON strain (*cmrRST*, CDR1689, CDR1690, CDR1914, and *cwp28*) were also more highly expressed in rough colonies compared to smooth, suggesting a role for these genes in colony morphology development. The 11 genes with higher transcript abundance in *cmr*Δ3-OFF compared to *cmr*Δ3-ON include *cwpV*, which encodes a cell wall protein, two uncharacterized genes (CDR0921 and CDR0922), and one gene encoding a spore coat protein (CDR2290) (18–20). The remaining seven genes are involved in ethanolamine utilization (CDR1834-1836, CDR1838-1841), four of which also had higher transcript abundance in smooth colonies compared to rough (CDR1834-1836, CDR1838). Overall, eleven of the 18 DEGs in the *cmr*Δ3-ON versus -OFF analysis were also differentially expressed in rough versus smooth isolates supporting a prominent role for *cmr* switch orientation in the transcriptional differences between rough and smooth colonies. However, *cmr* switch orientation does not account for all the differences between the naturally arising variants given that over 30 of the 50 DEGs between rough and smooth colonies were unique to that RNA-Seq comparison.

Additional pairwise comparisons and principal component analysis illustrate the similarities between the smooth, *cmr*Δ3-OFF, and Δ*cmrR*Δ*cmrT* colony transcriptomes and between the rough and *cmr*Δ3-ON colonies (**Fig. S1**). Notably, the only genes differentially expressed between Δ*cmrR*Δ*cmrT* and either the smooth isolate or the *cmr*Δ3-OFF mutant were *cmrR* and *cmrT*, and phase variable gene *cwpV*, which can likely be attributed to stochastic inversion of the *cwpV* switch (**Fig. S1A-B**, **Table S5**) (5, 19). Comparing transcript abundance in *cmr*Δ3-ON versus Δ*cmrR*Δ*cmrT* revealed 37 DEGs (**Fig. S1D**, **Table S4**), including all 18 DEGs identified in the *cmr*Δ3-ON versus - OFF comparison. Similarly, transcriptomic differences between the rough isolate and the Δ*cmrR*Δ*cmrT* mutant encompassed all the DEGs between rough versus smooth colonies (**Fig. S1C, Table S4**). Collectively, these transcriptome analyses indicate that CmrRST accounts for many, but not all, transcriptional differences between rough and smooth colony variants despite the requirement for CmrRST in rough colony development.

### The CmrR and CmrT regulons overlap with rough and *cmr*-ON regulons

We next sought to identify genes regulated by CmrRST. The individual *cmrR* and *cmrT* genes were ectopically expressed (“pCmrR” and “pCmrT”) under the control of an anhydrotetracycline (ATc)-inducible promoter, and RNA-Seq was used to compare the transcriptomes to similarly grown vector controls. In the pCmrR versus vector comparison, 15 genes were differentially expressed and met a log_2_ fold change > 2 and FDR p-value < 0.01 cutoffs (**Fig. 2A**, **Table 1**). The four genes downregulated in the pCmrR strain encode a c-di-GMP phosphodiesterase (*pdcC*) (17, 21), an osmoprotectant transport system (CDR3074-3075), and a hypothetical protein (CDR1929). The eleven upregulated genes encode three proteins involved in amino acid metabolism (*cysA*, *cysM*, and CDR2932), two hypothetical proteins (CDR1689 and CDR1914), Cwp28, and five proteins involved in signaling (CDR2040, *pdcB*, and *cmrRST)*. We note that PdcB and PdcC undergo phase variation, so the observed differences in their expression are likely independent of CmrR/CmrT (21, 22). Increased transcript abundance of all three *cmr* genes in the pCmrR strain aligns with previous work establishing an autoregulatory role for CmrR (**Fig. 1A**) (7). Ten of the 15 DEGs in the pCmrR versus vector comparison were also differentially expressed between rough and smooth colonies, and six were differentially expressed between *cmr*Δ3-ON and *cmr*Δ3-OFF colonies.

**Figure 2.**
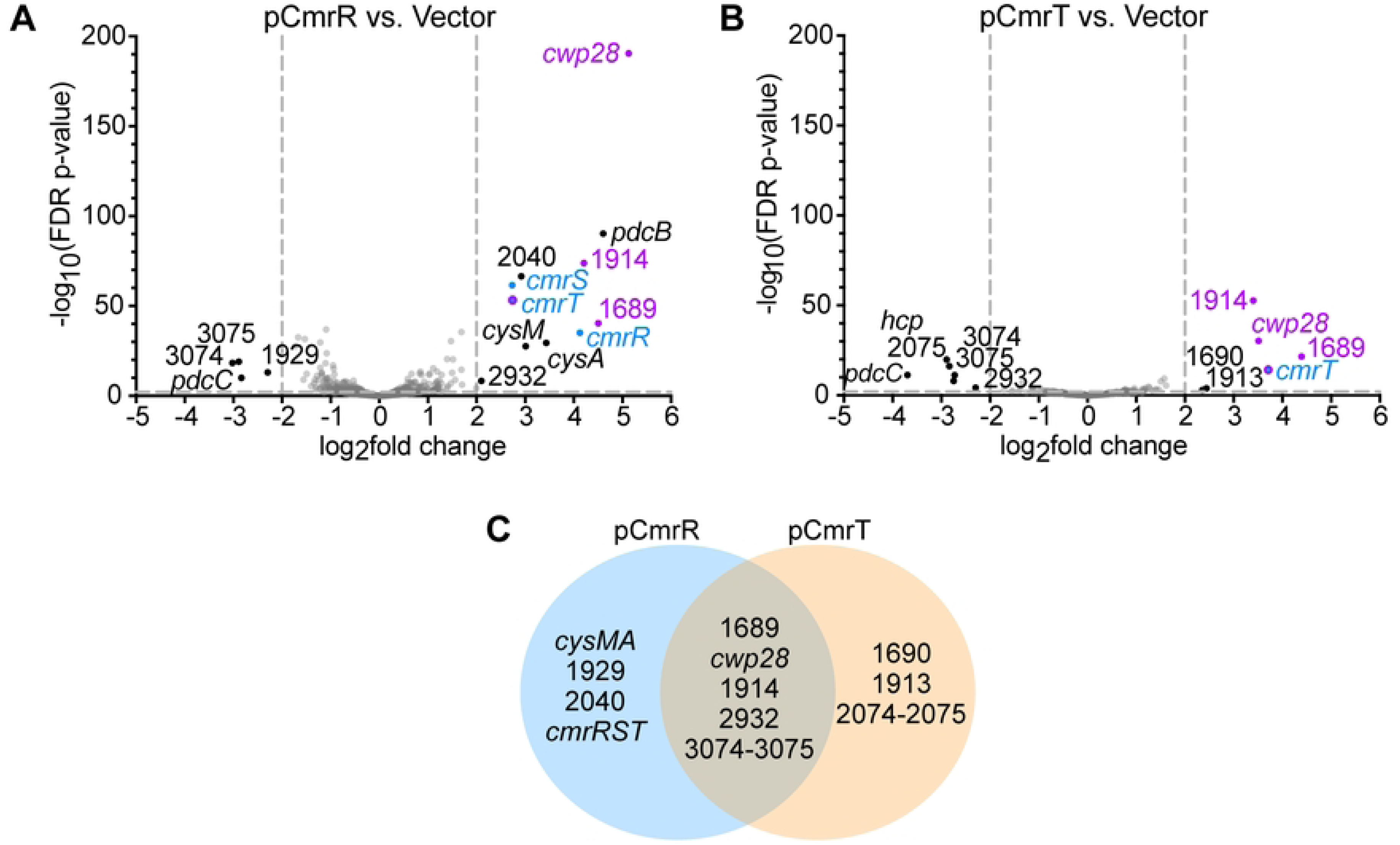
RNA-Seq identified genes regulated by CmrR and CmrT. **(A-B)** Volcano plots of differences in transcript abundances in strains grown in BHIS-Tm10-ATc broth. **(A)** Wildtype R20291 overexpressing *cmrR* (pCmrR) vs. the vector control. **(B)** Wildtype R20291 overexpressing *cmrT* (pCmrT) vs. the vector control. Gray dotted lines demarcate cutoffs of > 2 log_2_ fold-change and/or FDR *p*-value < 0.01. Black dots indicate genes differentially expressed genes that met cutoffs, gray dots indicate genes that did not meet the cutoffs, and blue dots denote *cmrRST*. Purple dots highlight genes that were differentially expressed in all four RNA-Seq comparisons (rough vs. smooth, *cmr*Δ3-ON vs. *cmr*Δ3-OFF, pCmrR vs. vector, pCmrT vs. vector). **(C)** Venn diagram of genes regulated by CmrR, CmrT, or both.

**Table 1.**
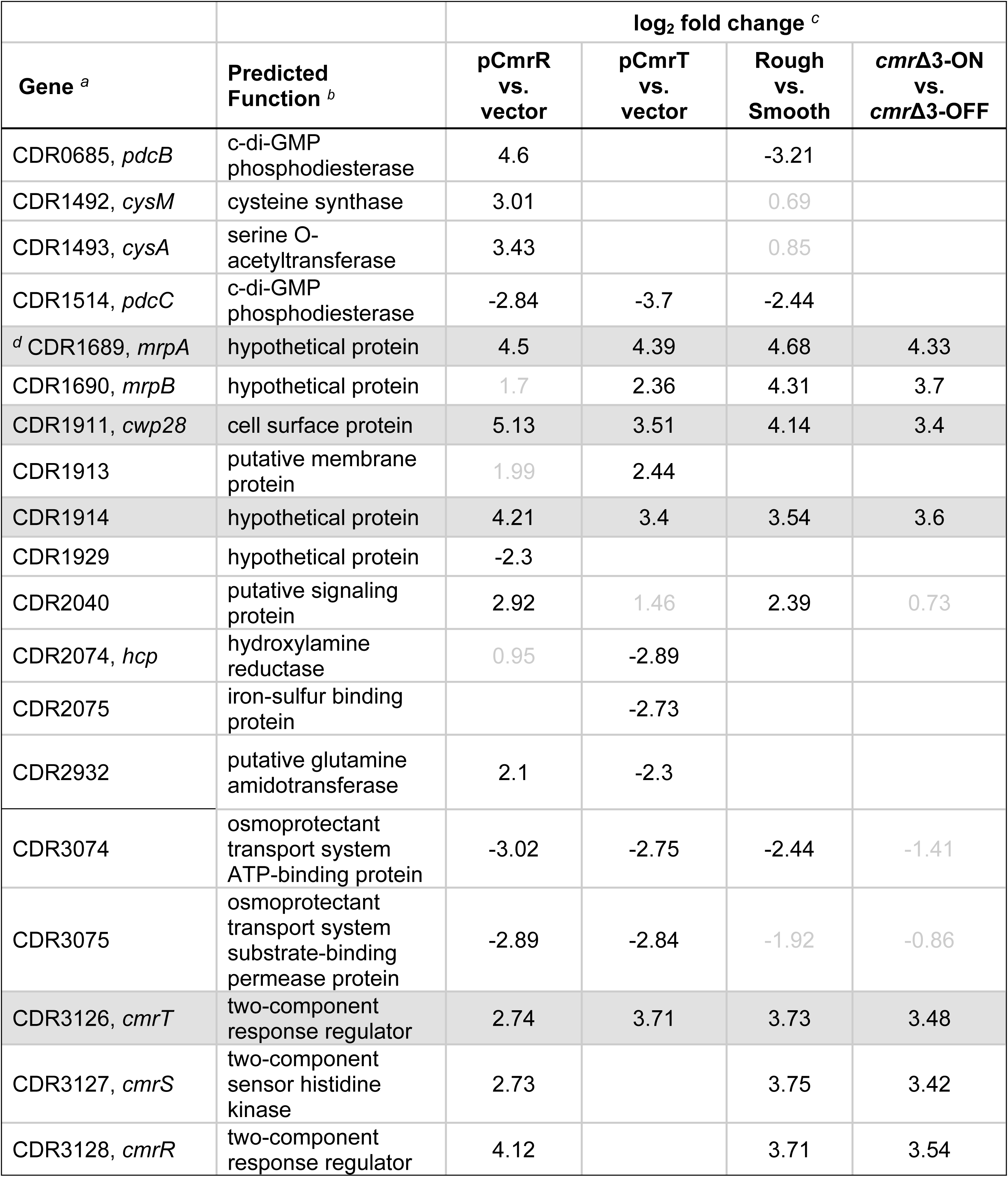
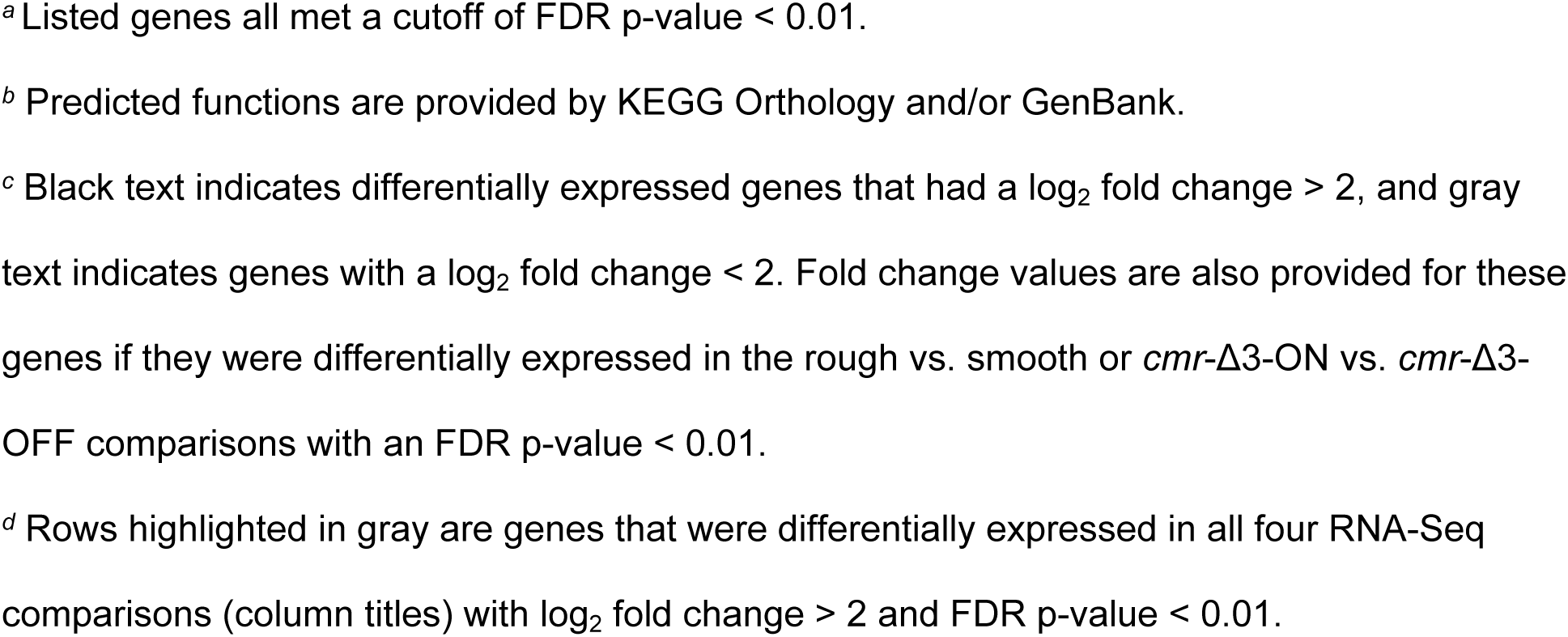
Genes differentially expressed with overexpression of *cmrR* or *cmrT* relative to the vector control strain.

Comparison of *C. difficile* pCmrT to the vector control yielded 12 genes meeting the fold-change and *p*-value cutoffs (**Fig. 2B**, **Table 1**). Six genes were downregulated by *cmrT* overexpression: the phosphodiesterase-encoding gene *pdcC,* three metabolism-related genes (CDR2074-2075 and CDR2932), and osmoprotectant transport system genes (CDR3074-3075). The six upregulated genes encode a putative membrane protein (CDR1913), Cwp28, three uncharacterized proteins (CDR1689, CDR1690, CDR1914), and CmrT itself. The elevated transcript abundance of *cmrT* (log_2_ fold change 3.71) confirms the successful overexpression of *cmrT*. Eight of the 12 CmrT-regulated genes were also differentially expressed between pCmrR and vector strains, indicating an overlap between CmrR and CmrT-regulated genes. Additionally, eight of the 12 CmrT-regulated genes were also differentially expressed between rough and smooth colonies, and five were differentially expressed between *cmr*Δ3-ON and *cmr*Δ3-OFF colonies.

In total, 63 genes were differentially expressed across all strains. Of note are four genes that were more highly expressed in the rough, *cmr*Δ3-ON, pCmrR, and pCmrT strains: *cmrT, cwp28,* and uncharacterized genes CDR1689 and CDR1914.

### CmrT is the predominant regulator of genes outside the *cmrRST* locus

Because prior work established an autoregulatory role for CmrR, but not CmrT (7), we assessed the contribution of each response regulator to differential expression of the genes identified by RNA-Seq. In a Δ*cmrR*Δ*cmrT* mutant, we individually expressed either *cmrR* or *cmrT* and used quantitative reverse-transcriptase PCR (qRT-PCR) to measure transcript abundance of genes that appeared to be regulated by CmrR only, CmrT only, or by both regulators (**Fig. 2C**, **Table 1**).

RNA-Seq analysis identified six genes as regulated by both CmrR and CmrT in the wildtype background (**Fig. 2C**, **Table 1**): CDR1689, *cwp28*, CDR1914, CDR2932, and CDR3074-3075. CDR2932 showed opposite regulation by CmrR and CmrT, while the other genes had similar trends with each regulator. Examination of the raw read count RNA-Seq data indicated that CDR2932 transcript abundance was low in all strains, so this gene was not investigated further. For CDR1689, *cwp28*, and CDR1914, expression of *cmrR* alone in the Δ*cmrR*Δ*cmrT* background did not alter transcript abundance, while *cmrT* expression alone resulted in significantly increased expression of all three genes (8.6-fold, 1.9-fold, and 9.2-fold, respectively) (**Fig. 3A-C**). CDR3075 was used as a proxy for both CDR3074-3075 because of a putative shared transcriptional start site upstream of CDR3075 (23). Although no significant differences in transcript abundance were observed when either *cmrR* or *cmrT* was expressed, transcript abundance was lowest when *cmrT* was expressed than in all other strains tested (**Fig. 3D**. These genes are thus regulated by CmrT, with indirect regulation by CmrR autoregulation of *cmrRST* transcription.

**Figure 3.**
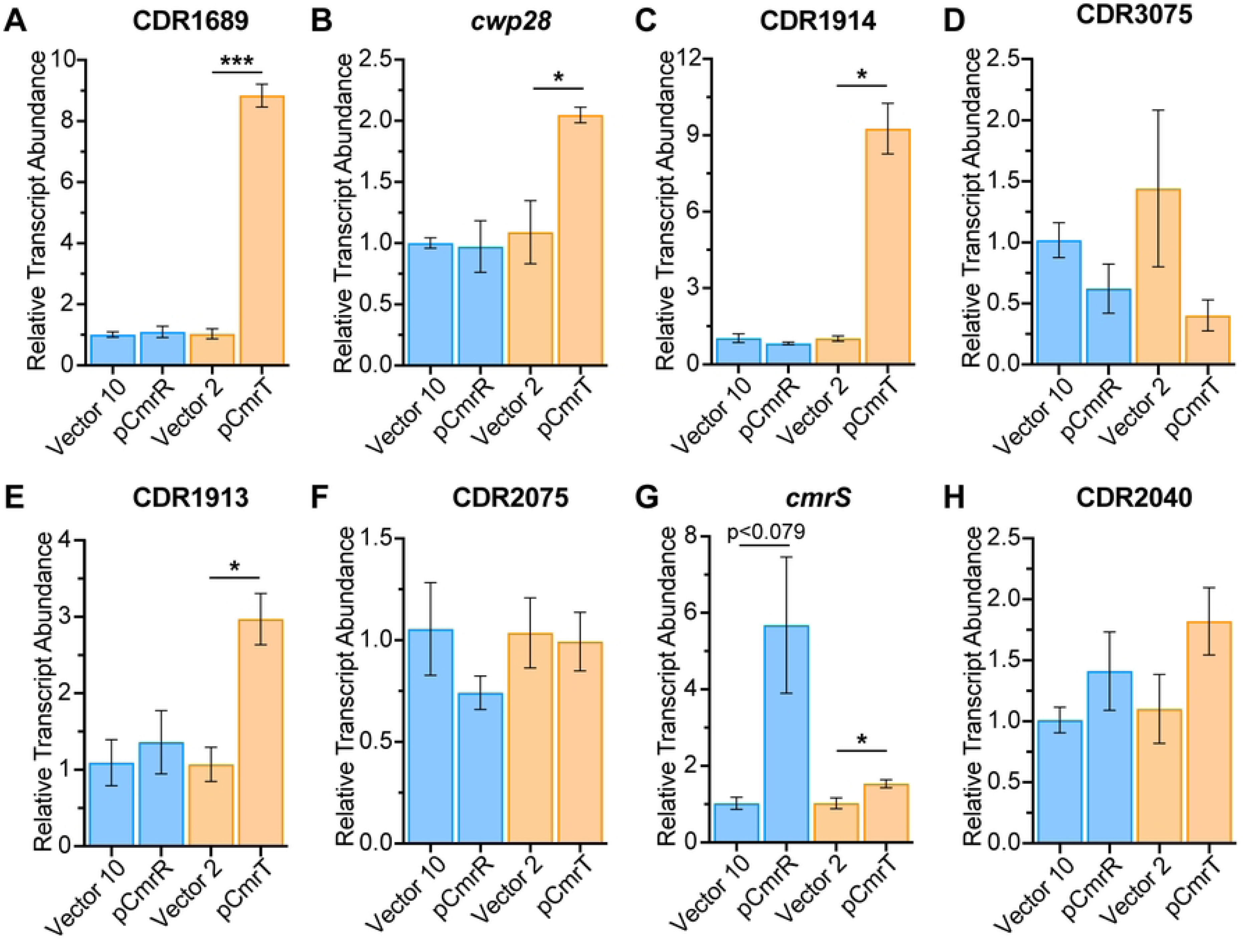
Relative contributions of CmrR and CmrT to gene regulation. **(A-H)** Relative transcript abundance determined by reverse-transcriptase qPCR for CDR1689 **(A)**, *cwp28* **(B)**, CDR1914 **(C)**, CDR3075 **(D)**, CDR1913 **(E)**, CDR2075 **(F)**, *cmrS* **(G)**, and CDR2040 **(H)** in Δ*cmrR*Δ*cmrT* ectopically expressing *cmrR* or *cmrT*. Cultures were induced with ATc at 10 ng/mL for pCmrR and the respective vector control (blue bars), and 2 ng/mL for pCmrT and the respective vector control (orange bars). Shown are means and standard error for three to four biological replicates. *p<0.05, ***p<0.001, unpaired two-tailed t-test with Welch’s post-test.

CDR1913 and CDR2075 were identified by RNA-Seq as genes exclusively regulated by CmrT. In the Δ*cmrR*Δ*cmrT* background, CDR1913 transcript abundance was significantly increased only when *cmrT* was expressed (2.8-fold), but not *cmrR* (**Fig. 3E**). We note that CDR1913 may also be indirectly regulated by CmrR, as RNA-Seq showed a significant difference in CDR1913 between pCmrR and vector, though this did not meet our cut-off for fold change. Neither *cmrR* nor *cmrT* expression significantly altered CDR2075 transcript abundance (**Fig. 3F**), suggesting both regulators are necessary for altered expression. CDR2040 and *cmrS* were exclusively regulated by CmrR in the RNA-Seq analysis. Consistent with previous work defining an autoregulatory role for CmrR (7), qRT-PCR showed ∼5.6-fold higher *cmrS* transcript upon the expression of *cmrR.* We also observed ∼1.5x higher *cmrS* transcript when *cmrT* was expressed (**Fig. 3G**). Neither CmrR nor CmrT alone was sufficient to alter transcript abundance of CDR2040 by qRT-PCR (**Fig. 3H**), suggesting roles for both regulators. Overall, these data indicate that most CmrRST-regulated genes require only CmrT for altered expression, *cmrRST* regulation occurs via CmrR, and a subset of genes may require both CmrR and CmrT for altered expression.

### CDR1689-1690, CDR1913-1914, and CDR1929 influence surface motility and rough colony development

To identify genes that mediate CmrRST-mediated rough colony development and surface motility on agar medium (3), we determined the impact of deleting or ectopically expressing CmrR- and CmrT-regulated genes on these two phenotypes. First, each of the CmrR- and CmrT-regulated genes highlighted in **Fig. 2C** was cloned for ATc-inducible expression (24); neighboring genes or genes that appear to be part of an operon were co-expressed. Because *cmrT* is required for rough colony development and surface motility, the genes were expressed in the Δ*cmrT* background to evaluate their sufficiency for these phenotypes (3). An inducible *cmrT*-expressing strain was included as a control. As seen previously, low-level induction of *cmrT* (2 ng/mL ATc) yielded rough colonies and increased surface motility, whereas higher inducer concentrations resulted in toxicity (**Fig. 4A-B**) (3).

**Figure 4:**
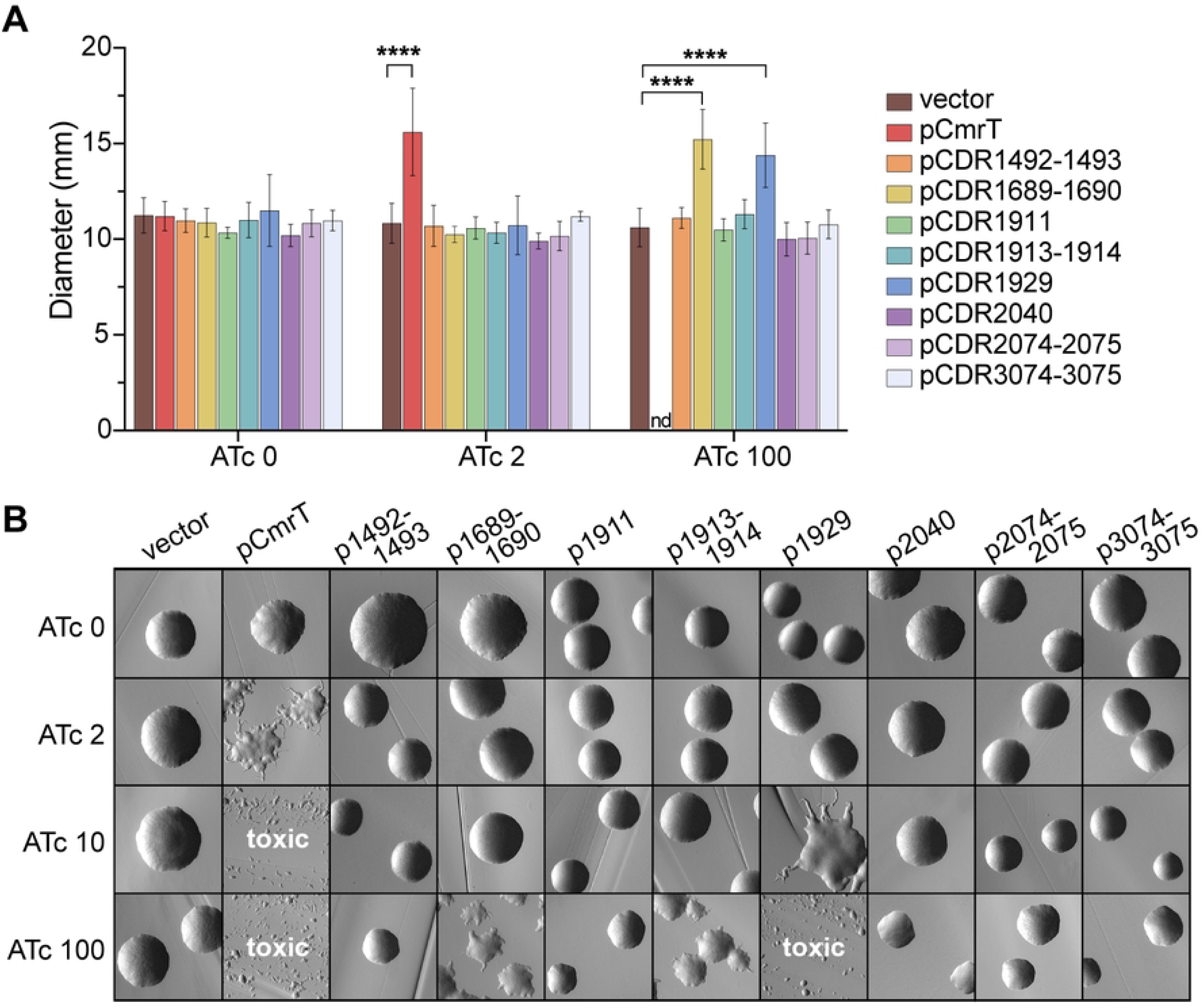
CDR1689-1690, CDR1913-1914, and CDR1929 promote surface motility and/or rough colony development. **(A)** Assay to determine the roles of CmrR- and/or CmrT-regulated genes in surface motility. Overnight cultures of Δ*cmrT* strains overexpressing CmrR- or CmrT-regulated genes or a vector control were spotted on BHIS-Tm10-agar and ATc 0-100 ng/mL. Surface motility was measured after seven days at 37°C. Shown are means and standard deviation for three biological replicates. ****p<0.0001, one-way ANOVA with Dunnett’s post-test. **(B)** Colony morphology of Δ*cmrT* strains overexpressing CmrR- and/or CmrT-regulated genes or the vector control grown on BHIS-Tm10-agar and ATc 0-100. Images were taken at 2X magnification after 24 hours at 37°C. Shown are representative images from four biological replicates.

While most strains resembled the vector control, two expression strains showed increased surface motility and a rough colony phenotype, and one strain had altered colony morphology with no discernable effect on surface motility. Specifically, expression of CDR1689-1690 with 100 ng/mL ATc restored surface motility and induced formation of rough colonies (**Fig. 4A-B**). CDR1929 expression with 100 ng/mL ATc significantly increased surface motility (**Fig. 4A**), but this ATc concentration resulted in sparse growth on agar medium (3) (**Fig. 4B, Fig. S2**). However, induction with a lower ATc concentration (10 ng/mL) yielded rough colonies (**Fig. 4B**). In contrast, ectopic expression of CDR1913-1914 did not affect surface motility with any ATc concentration but resulted in an intermediate colony phenotype with 100 ng/mL ATc (**Fig. 4A-B**).

To determine the requirement of the CmrRST-regulated genes for surface motility or rough colony phenotype, we successfully deleted CDR1689-1690, *cwp28*, CDR1913-1914, and CDR3074-3075. All the deletion mutants retained wildtype levels of surface motility, indicating that none are required for the phenotype (**Fig. S3A**). Additionally, all strains retained the ability to form both rough and smooth colonies (**Fig. S3A-B**). Therefore, while CDR1689-1690 are sufficient to promote surface motility and rough colony development, they are not required, suggesting that *C. difficile* has other mechanisms mediating these phenotypes. The role of CDR1929 could not be assessed because we were unsuccessful in creating a deletion mutant.

### Expression of genes that promote the rough colony morphology also decrease swimming motility

*C. difficile cmr*-ON variants have reduced swimming motility compared to *cmr*-OFF and Δ*cmrR*Δ*cmrT* (3, 7). To identify the CmrRST-regulated genes that inhibit swimming motility, we assessed the expression and deletion strains described above for swimming in soft agar medium. The Δ*cmrT* pCmrT strain, which exhibits reduced swimming motility, was included as a positive control (3). Ectopic expression of three sets of genes (CDR1689-1690, CDR1913-1914, and CDR1929) reduced swimming motility relative to the vector control, though not to the extent of pCmrT (**Fig. S4A**). These genes were determined to promote rough colony formation, suggesting a correlation between rough colonies and decreased swimming (**Fig. 4B**).

Swimming motility of the CDR1689-1690, *cwp28*, CDR1913-1914, and CDR3074-3075 deletion mutants were also assayed for swimming motility alongside *cmr*Δ3-ON and Δ*cmrR*Δ*cmrT* controls. After 48 hours of growth, swimming motility of all mutants was more similar to *cmr*Δ3-ON than to Δ*cmrR*Δ*cmrT*, indicating that these DEGs are not required for CmrRST-mediated inhibition of swimming motility (**Fig. S4B**).

### CDR1689-1690-mediated surface motility alleviates the selective advantage of the *cmr*-ON state

Prolonged incubation on an agar surface promotes *C. difficile* surface motility and selection of *cmr*-ON cells (3). Specifically, a *cmr*-OFF wildtype isolate grown on an agar surface exhibits surface motility and concurrent shift to a population with the *cmr* switch predominantly in the ON orientation. How selective pressures encountered during growth on an agar surface favor a *cmr*-ON population is unknown but presumably involves one or more CmrRST-regulated factors that enable surface motility. To explore this possibility, we first asked whether *cmr* switch inversion occurs during surface growth in the absence of surface motility. Wildtype *C. difficile* and a Δ*cmrR* mutant, which are capable of surface motility, and Δ*cmrT* and Δ*cmrR*Δ*cmrT* mutants, which lack surface motility, were grown on BHIS-agar for 3 days (3, 7). Quantitative PCR (qPCR) was then used to quantify the proportion of bacteria with the *cmr*-ON switch orientation before and after growth. The starting populations of all four strains were predominantly *cmr*-OFF (**Fig. 5A**). After 3 days of surface growth, the wildtype and Δ*cmrR* populations showed surface motility and a significant shift to *cmr*-ON, from ∼10% *cmr*-ON in the starting populations to 54% and 70% *cmr*-ON in the surface growth, respectively (**Fig. 5A**). These results indicate that CmrR is not required for phase variation to the *cmr*-ON state. In contrast, the Δ*cmrT* and Δ*cmrR*Δ*cmrT* populations remained in the *cmr*-OFF state (**Fig. 5A**), which suggests the phase variation to the *cmr*-ON state requires CmrT, and likely CmrT-dependent surface motility.

**Figure 5.**
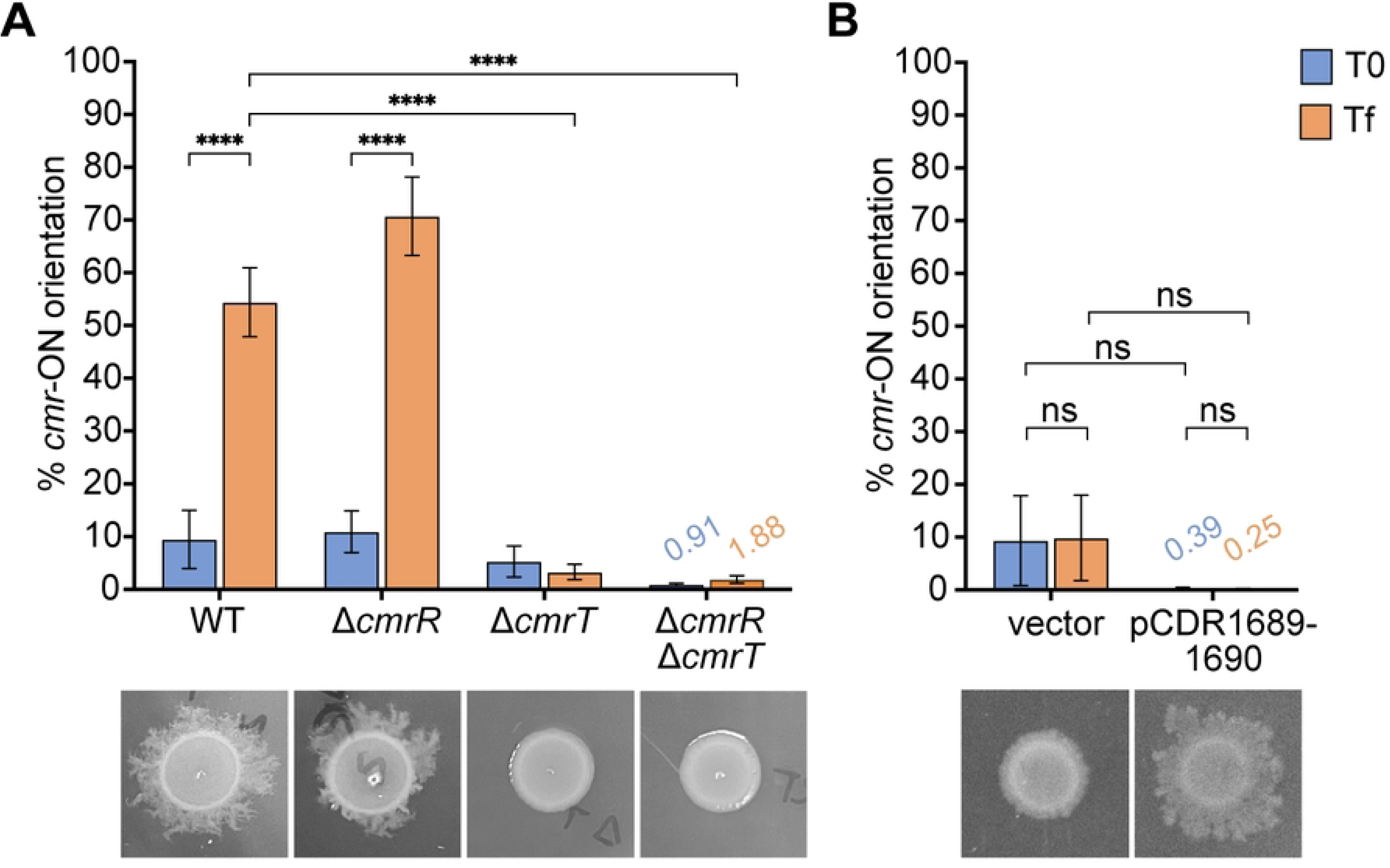
Surface motility conferred by CDR1689-1690 alleviates selection of *cmr*-ON cells. **(A)** qPCR was used to measure *cmr* switch orientation in WT, Δ*cmrR,* Δ*cmrT, or* Δ*cmrR*Δ*cmrT* populations before (T0, blue bars) and after (Tf, orange bars) three days of growth on BHIS-agar. **(B)** qPCR analysis of *cmr* switch orientation in the Δ*cmrT* mutant with vector or ectopically expressing CDR1689-1690 in the starting culture (T0, blue bars) and after seven days of growth on BHIS-Tm10-ATc100-agar (Tf, orange bars). **(A, B)** Below the x-axis are representative images of the strains after three **(A)** or seven **(B)** days of growth on BHIS-agar. Shown are the means and standard error for 4 – 5 biological replicates. Means are shown above bars as needed. ****p< 0.0001, two-way ANOVA with Tukey’s post-test. “ns”, not significant.

Based on these findings, we speculated that selection of *cmr*-ON cells depends on the expansion of that subpopulation on the agar surface. Because expression of CDR1689-1690 conferred surface motility in the absence of CmrT, we speculated that expression of these genes would relieve the selective pressures that favor the *cmr*-ON variant. To test this idea, we ectopically expressed CDR1689-1690 in the Δ*cmrT* background, then measured the proportion of cells with the *cmr*-ON switch orientation compared to a vector control before and after growth on BHIS-agar. While ectopic expression CDR1689-1690 restored surface motility as expected, the population remained *cmr*-OFF after seven days of growth (**Fig. 5B**). This result supports the conclusion that expression of CDR1689-1690, and presumably surface motility, provides *C. difficile* a sufficient advantage on an agar surface to alleviate selection of *cmr*-ON cells.

### CDR1689 and CDR1690 are small proteins that are highly conserved in *C. difficile*

The role of CDR1689-1690 in several *cmr*-associated phenotypes led us to characterize these genes further, first by determining the contribution of the individual genes to surface motility and rough colony development. When ectopically expressed individually in the Δ*cmrT* background, neither CDR1689 nor CDR1690 were sufficient to confer surface motility (**Fig. S5A**) or rough colony morphology (**Fig. S5B**), suggesting that both of these genes work coordinately to produce phenotypes associated with the *cmr*-ON state. Consistent with this result, we detected a transcript that spanned the length of the two genes indicating that CDR1689 and CDR1690 can be co-transcribed (**Fig. S5C**).

CDR1689 and CDR1690 are annotated as hypothetical proteins of 81 and 87 amino acids, respectively. To confirm that CDR1689 and CDR1690 are translated into proteins, we constructed C-terminal FLAG-tag translational fusions for CDR1689 and CDR1690, then expressed them individually in *E. coli* DH5⍺ and assessed translation via western blotting. For both CDR1689 and CDR1690, appropriately sized bands based on predicted molecular weight were detected, and the abundance of the proteins correlated with ATc concentration (**Fig. 6A**). Three-dimensional modeling with ColabFold (25) predicts the structures of CDR1689 and CDR1690 with high confidence using a per-residue confidence metric (predicted local distance difference test (plDTT) of > 90). CDR1689 is predicted to consist of five antiparallel β-sheets and one ⍺-helix towards the C-terminus (**Fig. 6B**). CDR1690 is predicted to contain four antiparallel β-sheets on one face of the protein, three antiparallel β-sheets on the other face, and one ⍺-helix (**Fig. 6C**). Because CDR1689 and CDR1690 are both required to restore motility and rough colony morphology in the Δ*cmrT* mutant (**Fig. S5A-B**), we also used ColabFold to predict the likelihood of heterodimer formation. Dimer modeling did not return a high confidence score where the two proteins are predicted to interact (plDTT ∼70), suggesting low likelihood of heterodimer formation (**Fig. S6A**). This prediction does not, however, account for multimers or additional protein partners that could influence protein conformation.

**Figure 6:**
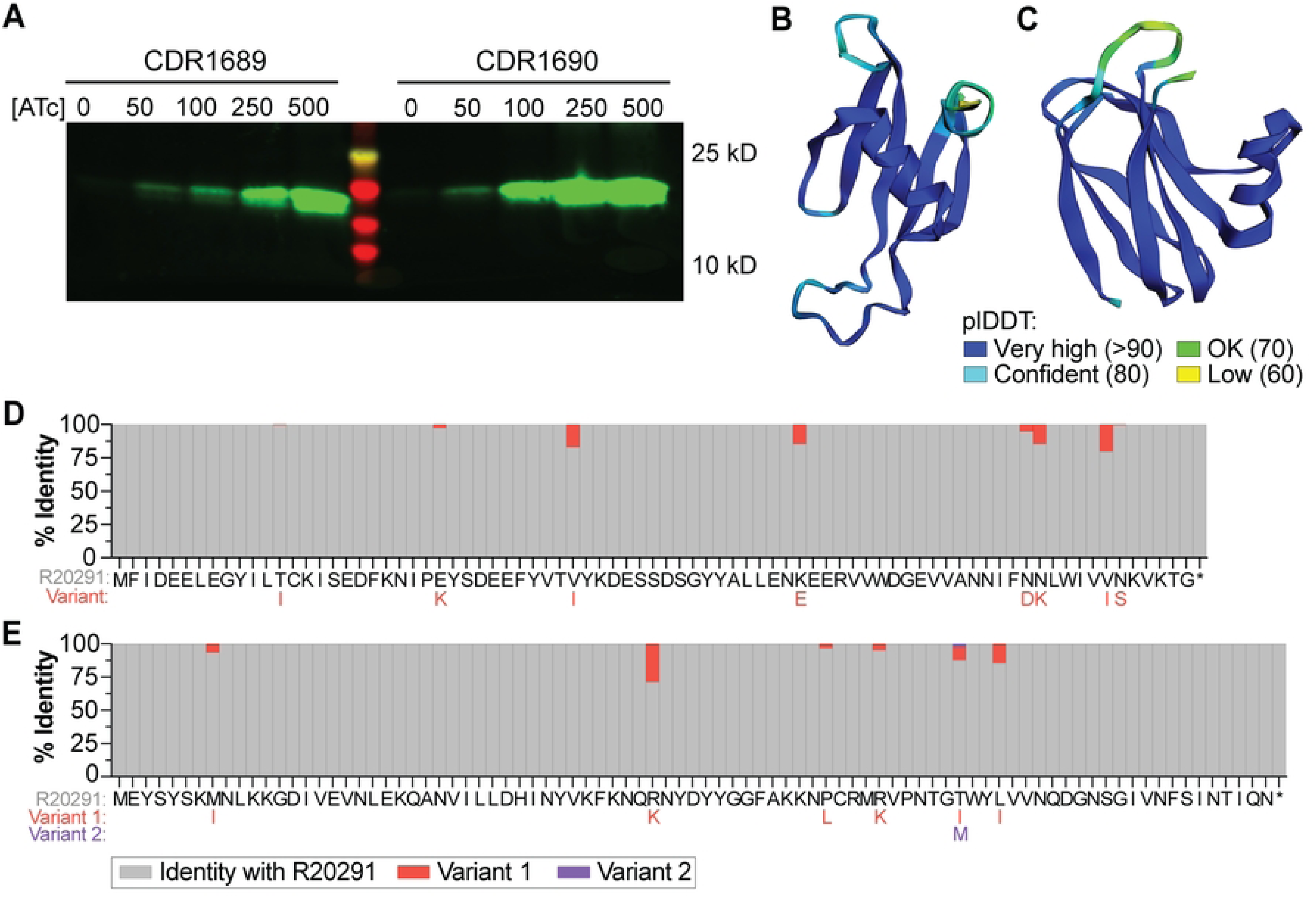
CDR1689 and CDR1690 encode small, uncharacterized proteins and are conserved in *C. difficile*. **(A)** FLAG-tag translational fusions of CDR1689 and CDR1690 were ectopically expressed in *E. coli* DH5⍺ cells with ATc 0-500 ng/mL, as indicated. Translated proteins were detected via western blot with anti-FLAG antibodies. CDR1689-FLAG is predicted to be 13.9 kilodaltons; CDR1690-FLAG, 14.5 kilodaltons. **(B, C)** Protein structures of CDR1689 **(B)** and CDR1690 **(C)** as predicted by ColabFold. Colors display the plDDT per-residue confidence metric indicated in the legend (bottom). **(D, E)** Amino acid conservation for CDR1689 **(D)** and CDR1690 **(E)** was determined across the 212 complete *C. difficile* genomes available on NCBI with RefSeq annotations. Along the x-axis are the amino acid sequences for each protein. Gray bars indicate the percentage of sequences that had the same amino acid as *C. difficile* R20291. Red and purple bars indicate the percentage of genomes that had a variant amino acid residue at that site; the substitutions are shown below the R20291 sequence in the respective red or purple text.

NCBI BLASTp revealed that both proteins are relatively restricted to *C. difficile*, with orthologs present in a limited number of closely related, unclassified *Clostridioides*, and the CDR1690 DUF1883 domain appearing in a range of bacterial genera (9, 26). To evaluate conservation of CDR1689 and CDR1690 across *C. difficile* strains, we searched for these amino acid sequences in the 212 complete genomes available on NCBI with RefSeq annotations. Both proteins were highly conserved in all 212 genomes that included several ribotypes across all five classical *C. difficile* clades, and the most mutable residues had conservative substitutions **(Fig. 6E-F)**. We named CDR1689 and CDR1690 <u>m</u>odulators of rough <u>p</u>henotype, *mrpA* and *mrpB*, respectively.

### MrpA interacts with MinD, a septum-site determining protein

MrpA and MrpB influence *cmr*-associated phenotypes through an unknown mechanism. While MrpA has no predicted protein family motifs, MrpB has three: a domain of unknown function (DUF1883) that spans the majority of the length of the protein, an overlapping bacterial pre-peptidase C-terminal domain (PPC), and a short Src homology 3 domain (SH3_2) (9). DUF1883 and PPC domains have similar predicted functions, with the DUF1883 domain predicted to be a ligand-binding domain of a PPC-like protein (27). In eukaryotes, SH3 domains are involved in protein-protein interactions that regulate the cytoskeleton (28, 29). Bacterial SH3 domains have been shown to mediate protein-protein and protein-peptidoglycan interactions (30–32). Given the wide range of potential functions by these domains and small size of MrpA and MrpB, we postulated that MrpAB mediate *C. difficile* phenotypes by interacting with other proteins. To identify potential interacting partners, we used affinity purification and mass spectrometry with the C-terminally FLAG-tagged MrpA or MrpB as bait. The MrpA-FLAG and MrpB-FLAG translational fusions were expressed in a *C. difficile* rough colony variant background to increase the chances of capturing protein-protein interactions that occur in rough or *cmr*-ON colonies. Following immunoprecipitation and liquid chromatography-tandem mass spectrometry, enriched proteins were identified by comparing the peptides associated with the FLAG-tagged proteins to those in the vector control.

Pulldowns with MrpA identified 258 proteins meeting the criteria of log_2_ fold change ≥ 1, q-value < 0.05, and sequence coverage > 50% (**Table S6**). As expected, MrpA was the most abundantly detected protein, confirming successful overproduction of the bait protein. pMrpB-FLAG pulled down 197 proteins meeting the above criteria, with MrpB being the most abundant (**Table S7**). Notably, MrpB pulled down CDR1689 with a log_2_ fold change of 3.45 over the vector control. The reciprocal interaction was not observed; it is possible that the C-terminal FLAG-tag on MrpA interfered with its ability to pull down MrpB. Overall, 117 proteins uniquely interacted with MrpA, 56 proteins were unique to MrpB, and 141 proteins were found to interact with both MrpA and MrpB.

The most enriched proteins for MrpA are the septum-site determining protein MinD (CDR0987, DivIVB), a probable glycine dehydrogenase (CDR1556, GcvPB), and a phosphotransferase system (PTS) component (CDR2780, LicB) (**Table S6**). MrpB also interacted with MinD and GcvPB, though the proteins most enriched by MrpB were an uncharacterized protein (CDR2123) and two components of a PTS (CDR2902 and 2901), followed by MrpA (**Table S7**). Given that CmrRST promotes cell elongation and chaining (3), we became interested in the interaction of MrpA with MinD. Of note, multiple other proteins involved in cell shape or structure were enriched by both MrpA and MrpB, including septum-site determining protein MinC, S-layer protein SlpA, cell-shape determining protein MreB, peptidoglycan-crosslinking enzyme Cwp22, and cell-division initiation protein DivIVA (CDR2503). MrpA also interacted with cell division protein FtsZ. These findings suggest a role for MrpA and MrpB in cell morphology or division. None of the identified proteins were identified in the RNA-Seq analyses.

To validate the interaction of MrpA and MrpB with MinD, we used a bacterial two-hybrid system in *E. coli* (33, 34). We created N- and C-terminal fusions of MrpA, MrpB, and MinD to the T18 and T25 subunits of the adenylate cyclase CyaA, then examined pairwise interactions via CyaA-dependent β-galactosidase activity. Both N- and C-terminal fusions of MrpA interacted with the respective MinD fusions (**Fig. 7A, Fig. S6B-C**). We were unable to confirm interactions between MrpA and MrpB or between MrpB and MinD, nor did we detect homodimerization of MrpA, MrpB, or MinD (**Fig. 7A, Fig. S6B-C**). These negative results are inconclusive, since the fusion of the large CyaA subunits with these small *C. difficile* proteins may interfere with proper protein folding or binding, the proteins may not interact directly, or *E. coli* cell division protein homologs may compete for binding. Together, the pulldown experiments and bacterial two-hybrid assay reveal an interaction between MrpA and MinD.

**Figure 7.**
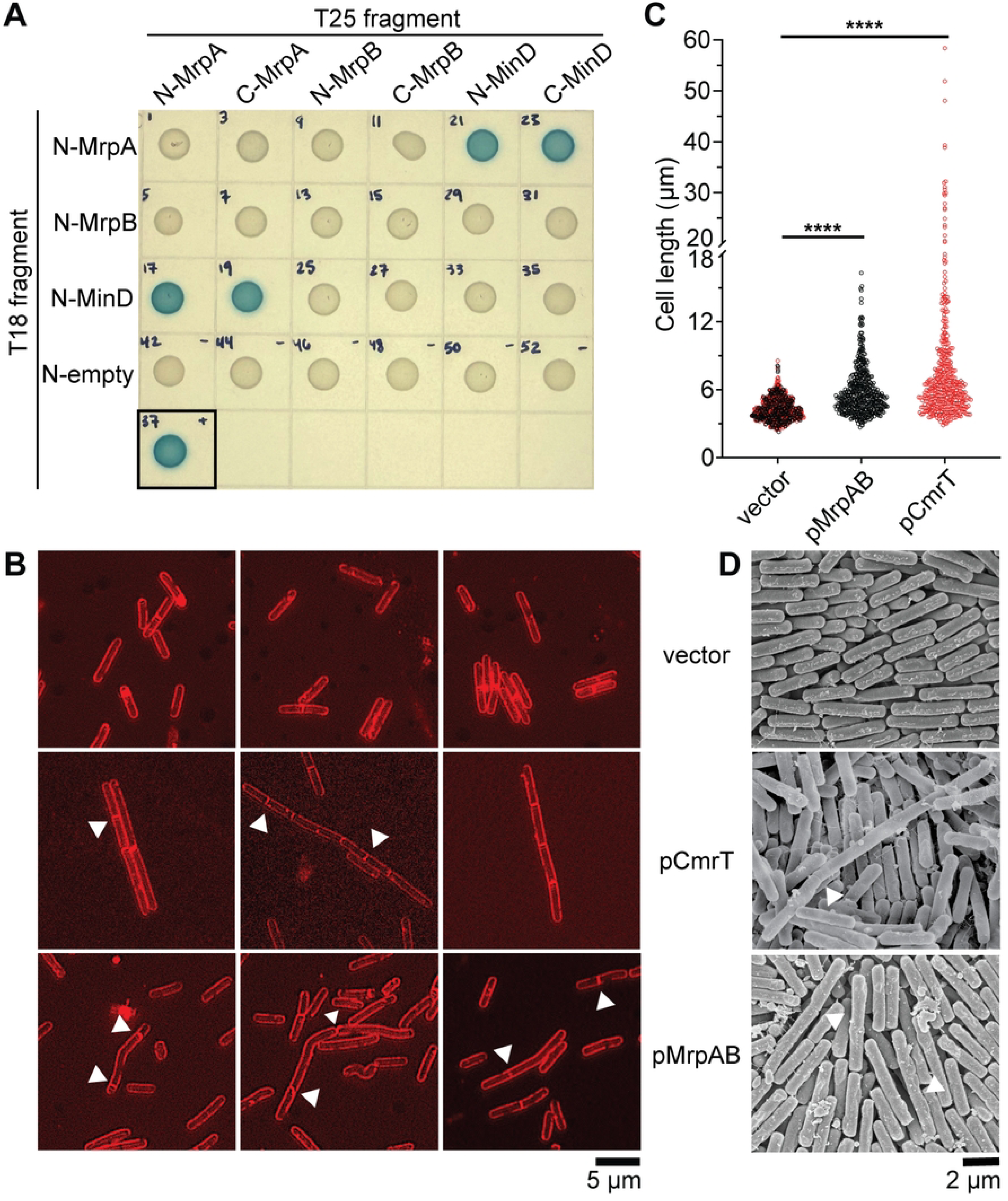
MrpA interacts with MinD. **(A)** Bacterial two-hybrid assay in which CDR1689, CDR1690, and MinD were translationally fused to the N-terminus of the T18 CyaA fragment and tested for interactions with CDR1689, CDR1690, or MinD fused to the N- or C-terminus of the CyaA T25 fragment. Overnight cultures of *E. coli* BTH101 carrying the indicated plasmid pair were spotted on LB-Kan50-Amp100-agar supplemented with 40 µg/mL X-gal and 0.5 mM IPTG. A T18 fragment with no translational fusion (“empty”) is included as a negative control. The black box indicates the zip-zip positive control. **(B)** FM™ 4-64 was used to stain membranes of Δ*cmrT* cells ectopically expressing CDR1689-1690, *cmrT*, or a vector control. Overnight cultures were grown in TY-Tm10 broth supplemented with ATc 2 ng/mL (vector and pCmrT) or ATc 100 ng/mL (vector and p1689-1690). Cells were imaged at 100X. Shown are representative images from two biological replicates. White arrows indicate asymmetric septa. **(C)** Quantification of cell lengths from FM™ 4-64 staining. Cells were measured from at least 10 different images from two biological replicates. Vectors induced with ATc 2 and ATc 100 ng/mL were combined; vector *n* = 359 (ATc 2 ng/mL, red dots) and *n* = 262 (ATc 100 ng/mL, black dots), pMrpAB *n* = 478, pCmrT *n* = 444). ****p< 0.0001, one-way ANOVA with Dunnett’s post-test. **(D)** Scanning electron microscopy images of Δ*cmrT* ectopically expressing CDR1689-1690, *cmrT*, or a vector control grown on BHIS-Tm10-ATc100-agar for 4 days. White arrows indicate asymmetric septa. Cells were imaged at 10,000X.

To determine if the MrpA-MinD interaction affects *C. difficile* cell division, we used membrane stain FM™ 4-64 to visualize cell morphology and septum formation in Δ*cmrT* cells ectopically expressing *mrpAB*, *cmrT*, or carrying vector controls. We found that *mrpAB* expression led to elongated cells, though fewer chained cells appeared compared to the pCmrT positive control **(Fig. 7B-C)**. Atypical cell division was apparent in both the pCmrT and pMrpAB strains; asymmetric or polar septa were present in both strains, and multiple septa also occurred, unlike in the vector control. These data corroborate the pulldowns and bacterial-two hybrid assays by providing phenotypic evidence that MrpA, and potentially MrpB, interact with MinD to disrupt *C. difficile* cell division.

## DISCUSSION

*C. difficile* colony morphology and multiple additional associated phenotypes undergo phase variation via the ON/OFF switching of CmrRST production. In this study, we used RNA-Seq analysis to characterize the transcriptional differences between the rough and smooth colony variants, determined the contribution of CmrRST to these differences, and identified two CmrRST-regulated genes, *mrpAB*, that promote the rough colony formation and surface motility.

Our multi-pronged RNA-Seq study revealed that CmrRST accounts for many, but not all, transcriptional differences between rough and smooth colony variants. Specifically, our results suggest that CmrRST activity drives development of rough colonies. Targeted analysis of these transcripts by qRT-PCR indicated that CmrT is the more immediate regulator of genes outside the *cmrRST* locus. For instance, expression of *cmrT* in the absence of *cmrR* was sufficient to alter *mrpA*, *cwp28,* CDR1913, and CDR1914 transcription. Whether CmrT directly activates transcription remains unknown, and there is no obvious consensus binding site upstream these genes.

Expression of *cmrR* alone did not significantly affect transcription of any genes other than *cmrRST*. CmrR may strictly serve an autoregulatory role, indirectly regulating genes by controlling *cmrT* expression (7). Because CmrT is a pseudoreceiver containing a glutamic acid residue at the usual response regulator phosphorylation site, modulation of CmrT levels may be sufficient to mediate *cmr*-associated phenotypes (3, 7, 35). It remains possible that CmrR works in conjunction with CmrT for a subset of genes: expression of *cmrR* or *cmrT* individually was insufficient to alter expression of at least three genes (CDR2040, CDR2075, and CDR3075). We speculate that CmrR and CmrT may form heterodimers to control expression of a subset of genes, or there may be a more complex regulatory relationship between these two proteins and the CmrS sensor kinase (3, 7).

Multiple genes were consistently associated with the rough colony/*cmr*-ON state, including *mrpAB*, *cwp28*, CDR1914, and CDR3074-3075. Through over-expression, we found that *mrpAB* and CDR1929 expression in a Δ*cmrT* mutant restored both rough colony formation and surface motility while inhibiting swimming motility. While CDR1929 was not further pursued due its toxicity when over-expressed, *mrpAB* was investigated for a role in CmrRST phase variation. We previously showed that growth of *C. difficile* on an agar surface selects for the *cmr*-ON variant (3). In the absence of *cmrT*, surface motility is lost along with selection of *cmr*-ON cells. Expression of *mrpAB* in the Δ*cmrT* mutant restored surface motility but not the enrichment of the *cmr*-ON variant. Thus, MrpAB-mediated surface motility alleviates selection of *cmr*-ON cells, suggesting that surface motility is necessary for expansion of the *cmr*-ON population.

Notably, none of the CmrRST-regulated genes were required for rough colony formation or surface motility. Redundancy is unlikely given the diverse gene functions. Our stringent criteria for differentially expressed genes may have excluded candidate genes. For example, *cwlA*, which encodes an endopeptidase necessary for cell separation (36, 37), showed ∼2-fold lower expression in the rough colony variant, the *cmr*Δ3-ON mutant, and the pCmrR strain compared to the respective smooth, *cmr*Δ3-OFF, or vector counterparts. Downregulation of *cwlA* could contribute to cell separation defects that result in chained cells in rough colonies and enhanced surface motility.

MrpA was found to interact with the septum site-determining protein MinD, revealing a potential mechanism for CmrRST-mediated phenotypes. The Min proteins are well characterized in *E. coli* and *B. subtilis* (11, 12, 38). In *E. coli*, MinE causes oscillation of MinD between cell poles, and MinD interacts with MinC, an inhibitor of FtsZ protofilaments. The accumulation of MinCDE at the poles help localize FtsZ to the mid-cell for septum formation. In *B. subtilis*, DivIVA localizes MinCD to cell poles to create a stationary gradient that promotes FtsZ polymerization at the mid-cell (12, 13, 39). *C. difficile* appears to have a hybrid of these two systems by encoding MinCDE and DivIVA, though the dynamics of these proteins are not well established (40, 41). In both *B. subtilis* and *E. coli*, *minD* mutants undergo abnormal cell division by forming both elongated and minicells (42, 43). Notably, a *B. subtilis minCD* mutant displays asymmetric, asynchronous cell division that results in elongated cells with irregularly spaced septa (44). We observed a similar phenotype in *C. difficile* expressing *mrpAB*. We postulate that the MrpA-MinD interaction disrupts typical cell division to result in the observed elongated, asymmetrically dividing cells. MrpA may reduce MinD activity or interfere with interactions with other cell division proteins.

The role of MrpB is unclear. Its co-expression with MrpA is required to elicit rough colony development and surface motility. Pulldown experiments indicated that MrpA and MrpB interact directly, though this latter result could not be corroborated with a bacterial two-hybrid assay. MrpB contains a bacterial SH3 domain, which frequently targets cell wall motifs and is often found fused to cell wall hydrolases (30–32), and a pre-peptidase domain. MrpB may thus be part of a larger complex influencing cell division. Supporting this hypothesis, we found that both MrpA and MrpB interacted with several proteins involved in cell shape and cell division that are already known to interact, including MinD, MinC, DivIVA, and FtsZ (38, 40, 41, 45).

Based on these and previous results, we propose that the elongated, chained cells in the *cmr*-ON rough variant contribute to surface migration as non-separated cells are pushed along the axis of cell division and growth (3). This process resembles sliding motility, in which expansion of a colony is driven by cell division and population growth (46–52). In *B. subtilis*, sliding motility is the result of differentiated cell types; surfactin-producing cells facilitate colony expansion by matrix-producing cells that form long, filamentous loops called “van Gogh bundles” that exhibit sliding motility and have impaired swimming motility (48). In *C. difficile*, inhibition of swimming motility may similarly be an indirect consequence as daughter cells remain tethered to mother cells, physically impeding flagellum-dependent motility. While no surfactant has been identified in *C. difficile* thus far, some bacteria exhibit sliding motility via alternate mechanisms. In *Salmonella enterica* serovar Typhimurium, sliding motility is thought to be facilitated by a surface protein (49, 53). *Mycobacterium smegmatis*, which also displays colony dimorphism correlated with motility and virulence, appears to rely on glycopeptidolipids for sliding motility (46, 54, 55).

Why certain environments favor the *cmr*-ON or *cmr*-OFF states is unknown. Presumably, the ability to migrate via different mechanisms allows access to nutrients (or avoidance of repellants), conferring a fitness advantage to the subpopulation that is more motile in the given environment. Consistent with this idea, many of the genes upregulated in smooth colonies function in metabolism, and computational modeling predicted the variants have distinct metabolic needs (56). Specifically, the smooth colony variant, but not the rough, was predicted to rely on glucose via the pentose phosphate pathway—a prediction supported by experimental evidence that in the absence of glucose, *C. difficile* develops rough colonies. Ongoing studies will examine the contributions of CmrRST-regulated genes and the dynamics of *cmr* switch inversion during infection of a murine model of disease, which may elucidate the effects of *C. difficile* phase variation on its survival in a complex and changing intestinal environment.

## METHODS

### Bacterial strains and growth conditions

*C. difficile* R20291 strains were grown statically at 37°C in an anaerobic chamber (Coy Laboratories) with an atmosphere of 85% N_2_, 5% CO_2_, and 10% H_2_ (Airgas). *C. difficile* was grown from freezer stocks on BHIS (37 g/L Brain Heart Infusion (BD 211059), 5 g/L yeast extract (Gibco 212750), and 1 g/L cysteine) with 1.5% agar. Overnight cultures (16-18 hours of growth) were grown in TY broth (30 g/L tryptone, 20 g/L yeast extract, 1 g/L sodium thioglycolate) and subcultured as indicated for each assay below. Media were supplemented with 10 µg/mL thiamphenicol (Tm10) for plasmid maintenance as needed. Ectopic expression of genes from the pRPF185 vector was induced with anhydrotetracycline (ATc, 0–100 ng/mL, as indicated). *E. coli* was grown aerobically in Luria-Bertani (LB) broth (Fisher BP1426 for broth, Fisher BP1425 for agar) at 37°C (for DH5⍺ and HB101(pRK24)) or 30°C (for BTH101). Antibiotics were included in the media as appropriate at the following concentrations: 10–20 µg/mL chloramphenicol (Cm10 or Cm20) and 100 µg/mL ampicillin (Amp100).

Table S1 lists all strains used in this study, and oligonucleotides are in Table S2. Details on plasmid and strain construction can be found in Text S1. Sequence-verified plasmids were transformed into HB101(pRK24) for conjugation into *C. difficile* R20291 as previously described (57). *C. difficile* expression strains were confirmed by PCR, and deletion mutants were confirmed with sequencing.

### RNA-Seq

Rough and smooth colony isolates of wildtype R20291, *cmr*Δ3-ON and *cmr*Δ3-OFF mutants, and the Δ*cmrR*Δ*cmrT* mutant were spotted (10 µL) on BHIS-1.5% agar in triplicate. After 24-hours incubation at 37°C, the growth was collected in TRIzol for RNA extraction isopropanol precipitation as described previously (57, 58). Plasmid-bearing strains (pCmrR, pCmrT, or vector only) were grown from a single colony in TY-Tm10 broth overnight, then diluted 1:30 in BHIS-Tm broth. At an optical density at 600 nm (OD600) of ∼ 0.3, expression was induced with ATc. *C. difficile* with pCmrR or the vector control were induced with 10 ng/mL ATc; because higher expression of *cmrT* is toxic to *C. difficile* (3), the pCmrT and vector control strains were induced with 2 ng/mL ATc. Strains with pCmrR or pCmrT were grown in triplicate, and vector controls were grown in duplicate. At OD600 ∼1, 2 mL of each culture was collected for RNA purification. RNA was purified by TRIzol extraction and isopropanol precipitation (57, 58).

Purified RNA was submitted to Genewiz (Azenta Life Sciences) for paired-end sequencing with kits used per manufacturer’s instructions. Briefly, rRNA was depleted using the Ribo Zero rRNA Removal Kit (Illumina). RNA sequencing libraries were prepared with the NEBNext Ultra RNA library prep kit for Illumina (NEB) and checked using a Qubit 2.0 fluorometer. The libraries were multiplexed for sequencing using a 2 × 150 paired-end configuration on an Illumina HiSeq 2500 instrument. Image analysis and base calling were done using the HiSeq control software. The resulting raw sequence data files (.bcl) were converted to the FASTQ format and demultiplexed with bcl2fastq 2.17 software (Illumina). One mismatch was permitted for index sequence identification. Sequencing files are available on NCBI GEO (Accession: GSE305463).

Differential expression analysis was done as previously described using established bioinformatics tools (59). Briefly, Illumina reads were trimmed using Trimmomatic (60) then mapped to the *C. difficile* R20291 genome (GenBank: FN545816.1) using Bowtie2 (61). Mapped reads were then assigned to genes via FADU (62). DESeq2 analysis was used for pairwise comparison between the indicated strains (63).

### Quantitative reverse transcriptase PCR

Overnight cultures of *C. difficile* were diluted 1:30 in BHIS-Tm10 and induced with ATc (2 ng/mL for pCmrT and vector control, 10 ng/mL for pCmrR and vector control) at OD600 ∼0.3. At OD600 ∼1, 2 mL of culture was pelleted and stored in 1:1 ethanol:acetone at −80°C. RNA was extracted as described previously (57, 58). Purified RNA was reverse transcribed and quantitative reverse transcriptase PCR (qRT-PCR) was carried out as previously described with normalization to reference gene *rpoC* (3, 21, 64). No reverse transcriptase controls were run in parallel for all samples. Primer sequences can be found in Table S2.

### Orientation-specific qPCR

Overnight cultures were diluted 1:30 in 3 mL BHIS broth. At OD600 ∼1, 5 µL of each culture was spotted on three BHIS-1.5% agar plates; 2 mL of the inoculating cultures were collected for genomic DNA extraction as previously described (57). After three days, surface growth from three spots was collected and pooled, standardized by OD600 to collect roughly the same number of cells, and pelleted for genomic DNA extraction. For ectopic expression of CDR1689-1690 in the Δ*cmrT* mutant background, overnight cultures (5 µL) were spotted on BHIS-Tm10-ATc100-1.5% agar plates, and 2 mL of the inoculating culture were reserved for DNA isolation. After seven days, surface motility was measured, and each spot was collected independently for genomic DNA extraction.

All genomic DNA samples from the inoculating cultures and subsequent surface growth were assessed for *cmr* switch orientation with orientation-specific qPCR as described previously (3, 21). The percentage of the population with the *cmr* switch in the ON orientation was determined as described previously using *rpoA* as the reference gene (5). Primer sequences can be found in Table S2.

### Multiple sequence analysis for CDR1689 and CDR1690 conservation

All *Clostridioides difficile* genomes available on NCBI were filtered to include only complete genomes that were annotated by NCBI RefSeq. The resultant 212 genomes and fasta files for proteins CDR1689 and CDR1690 were downloaded. FaToTwoBit Version v377 was used to convert each of the downloaded fna files in the genome directories to a 2bit format. BLAT version 36×2 (65) was then used to identify the DNA sequences corresponding to those proteins by querying the 2bit genomes for the protein sequence contained in the fasta files. The protein-coding sequences for each genome were then extracted into another fasta file, with the start and end points corrected to accommodate the start and stop codons for each sequence. Reverse-stranded sequences were base-complemented and reversed. Mafft version 7.525 (66) was used to align the resulting base sequences. Expasy’s Translate Tool (67) was then used to identify the correct reading frame for the base sequences, and then Python Version 3.13.3 and Biopython Version 1.85 (68) were used to convert the aligned DNA sequences to amino acid sequences. R version 4.4.0 and GraphPad Prism were used to produce the conservation plots from the amino acid sequences.

### Affinity pulldowns

CDR1689 and CDR1690 with a C-terminus serine-glycine linker and 3X-FLAG tag were expressed in *C. difficile* using the pRPF185 tetracycline-inducible expression vector (**Table S1, Text S1**) (24). To enrich for the rough colony morphotype, which expresses *cmrRST* at higher levels (3), overnights of *C. difficile* harboring pCDR1689-FLAG, pCDR1690-FLAG, or the vector control were spotted onto BHIS-Tm10-agar. After seven days of incubation at 37°C, rough tendrils were isolated and used for affinity co-purification.

Overnight cultures grown in 5 mL TY-Tm10 were diluted 1:100 into 500 mL BHIS-Tm10. Expression was induced with 100 ng/mL ATc when cultures reached an OD600 ∼0.2-0.3. At OD600 ∼1, cells were collected by centrifugation, supernatants were discarded, and pellets were stored overnight at −80°C. Once thawed, cell pellets were washed three times with 10 mL 1x PBS (137 mM NaCl, 2.7 mM KCl, 8 mM Na_2_HPO_4_, 1.5 mM KH_2_PO_4_) to remove residual culture medium, with centrifugation between washes occurring at 4,000 rpm, 10 minutes, 4°C. Washed cells were suspended in 3 mL of TNG buffer (50 mM Tris pH 6.8, 150 mM NaCl, 10% glycerol) supplemented with Pierce Protease Inhibitor Tablets (Thomas Scientific A32953) then lysed by bead beating (4x for 1 minute, 5 minutes on ice between rounds). Lysates were centrifuged at 10,000 x g for 15 minutes at 4°C, then supernatants were transferred to Protein LoBind tubes. Protein concentrations of the lysates were measured using the Pierce™ BCA Protein Assay Kit (Thermo Scientific A55864). For each sample, 4 mg total protein was added to 50 µL of packed anti-FLAG M2 magnetic beads (Sigma M8823) equilibrated in TNG buffer. Samples were incubated on an end-over-end rotor at 4°C overnight. Beads were washed six times in 1x PBS, suspended in 70 µL PBS, then transferred to a new Protein LoBind tube. A 5% aliquot of each sample was saved for western blot analysis, and the remainder was stored at −80°C. Protein identification via LC-MC/MS is detailed in Text S1.

### Microscopy

For imaging whole colonies, strains were streaked from freezer stocks on BHIS-agar (and Tm as required) and incubated for one day. Isolated colonies were streaked again on BHIS-agar without 0.1% cysteine supplementation (with Tm as required). After 24 hours of incubation, colonies were imaged at 2X magnification on a BZ-X810 microscope (Keyence).

Microscopy to determine cell morphology was conducted on Δ*cmrT* vector, pCmrT, or pCDR1689-1690 strains grown overnight in TY-Tm10 broth supplemented with ATc 2 ng/mL (vector or pCmrT) or 100 ng/mL (vector and pCDR1689-1690). For membrane staining, overnight cultures were suspended with 1 µg/mL FM™ 4-64 dye (Invitrogen T3166) for 15-30 minutes then pipetted on microscope slides covered with agarose pads. Cells were imaged at 100X magnification on a BZ-X810 microscope (Keyence). Cell lengths were measured using ImageJ software.

Scanning electron microscopy is detailed in Text S1.

### Bacterial two-hybrid

The *mrpA* (CDR1689), *mrpB* (CDR1690), and *minD* (CDR0987) sequences were codon-optimized for translation in *E. coli* and synthesized by Genewiz. The DNA fragments were cloned into pKT25, pKNT25, pUT18, and pUT18C to obtain N- and C-terminal fusions to CyaA domains (Euromedex) (**Text S1, Table S1**) (33, 69). Pairs of plasmids to be tested were co-transformed into chemically competent *E. coli* BTH101. Overnight cultures (5 µL) of these of BTH101 strains were spotted on LB-agar supplemented with 40 µg/mL 5-bromo-4-chloro-3-indolyl-β-D-galactopyranoside (X-gal), 0.5 mM isopropyl β-D-thiogalactopyranoside (IPTG), Amp100, and Kan50. Plates were imaged using standard photography after 24-36 hours of growth at 30°C.

## ACKNOWLEDGEMENTS

This work was supported by grants R01-AI143638 and R21-AI186221 to R.T. This research is based in part upon work conducted using the UNC Proteomics Core Facility, which is supported in part by NCI Center Core Support Grant (2P30CA016086-45) to the UNC Lineberger Comprehensive Cancer Center. Pulldown sample preparation and LC-MS/MS analysis was conducted by Thomas Webb and Dr. Aurora Cabrera, and data analysis was done by Angie Mordant and Dr. Laura Herring. The UNC Bioinformatics and Analytics Research Collaborative determined the protein conservation of MrpAB. Dr. Corbin Jones, Dr. Austin Hepperla, and Tyler Interrante assisted Christopher J. Serody with the analysis.

Kristen White, Jillann Madren, and Victoria Madden at The Microscopy Services Laboratory, UNC Department of Pathology and Laboratory Medicine conducted the scanning electron microscopy. The core is supported in part by P30 CA016086 Cancer Center Core Support Grant to the UNC Lineberger Comprehensive Cancer Center.

The content of this manuscript is solely the responsibility of the authors and does not necessarily reflect the official views of the funding agency.

## AUTHOR CONTRIBUTIONS

**A.M.:** Conceptualization, formal analysis, investigation, methodology, visualization, writing – original draft, writing – review and editing

**E.M.G.:** Conceptualization, investigation, methodology, writing – review and editing

**C.J.S.:** Formal analysis, investigation, methodology

**R.T.:** Conceptualization, formal analysis, funding acquisition, investigation, methodology, project administration, supervision, visualization, writing – original draft, writing – review & editing

